# Effects of 5-aza-2’-deoxycytidine on human osteoarthritic chondrocytes

**DOI:** 10.1101/476069

**Authors:** Shirin Kadler, Özlem Vural, Luzia Reiners-Schramm, Roland Lauster, Mark Rosowski

## Abstract

**Background:** Given regenerative therapies, the utilization of primary human cells is desired and requested in the development of *in vitro* systems and disease models. After a few passages *in vitro*, all cells from the connective tissue end up in a similar fibroblastoid cell type marked by loss of the specific expression pattern. It is still under discussion whether different de-differentiated mesenchymal cells have similar or identical differentiation capacities *in vitro*.

**Methods:** Chondrocytes isolated from patients with late-stage osteoarthritis were cultured for several passages until de-differentiation was completed. The mRNA level of cartilage markers was investigated, and the adipogenic, osteogenic and chondrogenic differentiation capacity was examined. By adding 5-aza-2’-deoxycytidine (5-aza-dC) to the media, the influence of DNA methylation on the differentiation capacity was analyzed.

**Results:** The chondrocytes used in this work were not affected by the loss of specific gene expression upon cell culture. The mRNA levels of SOX5, SOX6, SOX9, aggrecan, and proteoglycan-4 remained unchanged. The underlying mechanisms of cartilage marker maintenance in osteoarthritic (OA) chondrocytes were investigated with a focus on the epigenetic modification by DNA methylation. The treatment of de-differentiated chondrocytes with the DNA methyltransferase inhibitor 5-aza-2’-deoxycytidine (5-aza-dC) displayed no appreciable impact on the observed maintenance of marker gene expression, while the chondrogenic differentiation capacity was compromised. On the other hand, the pre-cultivation with 5-aza-dC improved the osteogenesis and adipogenesis of OA chondrocytes. Contradictory to these effects, the DNA methylation levels were not reduced after treatment with 1 μM 5-aza-dC for four weeks.

**Conclusion:** Chondrocytes isolated from late-stage osteoarthritic patients represents a reliable cell source for *in vitro* studies as wells as disease models since the chondrogenic differentiation potential remains. 5-aza-2’-deoxycytidine could not further improve their chondrogenic potential.

## Background

In multicellular organisms, the loss of environmental signals occurring during injuries leads to a stepwise reprogramming or de-differentiation of cells. On the other hand, cell fates are stabilized by epigenetic modifications as found in histone tail marks or DNA methylations. In this context, tissue-dependent differentially methylated regions (T-DMR) can be identified in differentiated cells. These cell type specific sequences are de-methylated in the regarding tissue and associated with active histone marks, while in other tissues these sequences are transcriptionally inactivated by methylation [1]. Therefore, the methylation status of key regulatory sequences represents a crucial factor for differentiation processes as well as cell phenotype maintenance and should be taken into consideration for successful regenerative applications [2,3]. The enzymes responsible for the DNA methylation are encoded by the genes DNMT1, DNMT3a, and DNMT3b (DNA methyltransferases). However, for cellular development, the introduction, as well as the removal of DNA modifications, are necessary. So far, no *in vivo* mechanism of direct DNA de-methylation is described, nevertheless modified intermediates as 5-hydroxymethylcytosine, 5-formylcytosine and 5-carboxylcytosine have been found [4]. These further modifications of 5mC (5-methylcytosine) are catalyzed by the TET family (ten-eleven translocation).

Caused by the withdrawal of tissue specific microenvironmental cues, cells de-differentiate upon *in vitro* cell culture, reflecting the loss of tissue specificity. The de-differentiation process of isolated human chondrocytes affects the epigenetic pattern of the cells on DNA methylation and histone modification level [5,6]. Alterations in DNA methylation are also identified in comparative studies of healthy chondrocytes with cells from osteoarthritic tissues [7,8].

Osteoarthritis (OA) is the most common disease of the osteochondral unit. OA is characterized by loss of permanent cartilage, reduced joint spaces, osteophyte development, subchondral bone cyst formation, and sclerosis. To compensate for the loss of matrix integrity, clusters of proliferating chondrocytes emerge in early osteoarthritic cartilage. The proliferation of hitherto resting articular chondrocytes is initiated by changes in the environment of the pericellular matrix (PCM). The disruption of the connection between chondrocytes and their PCM compromise the HA-CD44 signaling (hyaluronan and its receptor) and results in the up-regulation of MMPs with a decreased survival of cells [9]. With disease progression, fissures in the cartilage ECM can be detected. The degradation leads to an increase in oxygen tension that further accelerates the process of tissue destruction. Subsequently, the subchondral bone layer is affected, marked by tissue mass reduction. Furthermore, the subjacent calcified layer expands into the articular zone, and tissue vascularization is induced in late OA stages. The disease does not only affect the osteochondral unit of the joint but also the ligaments and the synovium [10]. Pharmaceutical intervention is the only option for patients to reduce pain since OA is not curable. In late state OA, the replacement of the joint by surgery is the only option for patients. Tissue engineering applications provide a promising strategy to regenerate the damaged tissues. In these technologies, the de-differentiation process is reversed by the cultivation of de-differentiated cells or stem cells in appropriate conditions capable of inducing the desired differentiation [11]. The cultivation conditions are oriented towards the *in vivo* process of chondrocyte differentiation and cartilage development by endochondral ossification [12]. The process of endochondral ossification is initiated by condensation of mesenchymal stem cells following differentiation steps of proliferative chondrocytes, hypertrophic chondrocytes and subsequent bone formation [13]. In parallel, articular cartilage differentiation and joint formation are regulated in a precise orchestrated developmental process [14]. For both, the differentiation of articular chondrocytes and the differentiated cartilage tissue, sets of marker molecules are well described. These sets include transcription factors like SOX5 and SOX9, signaling molecules from FGF-, BMP- and WNT-family as well as matrix molecules like collagen type 2 and aggrecan [13,15].

The molecules 5-azacytosine (5-aza) and 5-aza-2’-deoxycytosine (5-aza-dC) were first synthesized in 1964 [16] and have FDA approval for myelodysplastic syndrome treatment since 2004 and 2006, respectively. This base is a modification of cytosine, replacing the carbon at position 5 by nitrogen. This small alteration inhibits the functionality of DNA methyltransferases (DNMTs) by forming an irreversible covalent bond between the DNMT and the 5-aza-dC incorporated DNA strand [17]. Hence, the incorporation of 5-aza-dC into the DNA leads to a decreasing amount of functional DNMTs in the nucleus. It is reported by Cameron, Bachman, Myöhänen, Herman, & Baylin, (1999) and many others that cell lines treated with 5-aza-dC led to a decrease in DNA methylation and expression of formerly repressed genes.

To assess the impact on regenerative applications, we analyzed the influence of a DNMT inhibitor on chondrocytes isolated from patients with late-stage OA. The differentiation potential towards adipogenic, osteogenic and chondrogenic lineages was tested, and DNA methylation levels for specific target sites were analyzed.

## Methods

### Cell culture

Cartilage was extracted from hip and knee replacements surgeries of OA patients. Samples were isolated from non-weight bearing sections with no visible lesions. Human chondrocytes are the sole cellular component in articular cartilage. Therefore, no sorting strategy was necessary. The extracted cartilage was washed twice with PBS and shortly with 80% (v/v) ethanol followed by PBS and dissected from the underlying bone. The tissue was cut in small 2 mm x 2 mm pieces using a scalpel.

The pieces were digested with 1 mg/ml protease K (Sigma Aldrich) for 30 min, washed in PBS and digested in collagenase (2 mg/ml) overnight. A 70 μM cell strainer was used to remove the extracellular matrix debris from the cells. Chondrocytes were counted and 5 × 10^5^ cells were lysed for nucleic acid isolation. All remaining cells were seeded in a T25 cell culture flask for adherent growth using DMEM with 10 % FBS and 1% penicillin/streptomycin. Chondrocytes were passaged at 80% confluence. For experiments cells were used at passage 4 to passage 13. Even at late passages, no indications of senescence were found.

### AZA treatment

Non-confluent adherent cells were treated with 0.5 μM to 10 μM 5-aza-dC (100 mM stock solution in 25% acetic acid) in growth medium for a minimum of 3-4 weeks. Fresh 5-aza-dC was added every 24h, while the complete medium exchange was performed every other day.

### Osteogenic differentiation

Cells were seeded in 6-well plates (2.5 × 10^5^/well) and media was changed to osteoblast inducing conditions by adding 10 mM beta-glycerophosphate, 10 nM cholecalciferol (vitamin D3), 100 μM ascorbate phosphate and 10 mM dexamethasone in final concentrations for 28 days. The medium was changed every other day. Differentiation was visualized by Alizarin Red staining and confirmed with qPCR by testing mRNA level for osteopontin (OPN).

### Alizarin Red staining

The medium was carefully aspirated from cell culture wells and washed twice with dH_2_O. Alizarin Red solution (2% in dH_2_O at pH 4.1-4.3) was applied to the wells and incubated at room temperature for 5-10 minutes. The staining solution was removed carefully, and wells were gently washed twice with dH_2_O. Pictures were taken immediately.

### Adipogenic differentiation

Cells were seeded in 6-well plates (2.5 × 10^5^/well), and media was changed to adipogenic inducing conditions by adding 10 μg/ml insulin, 500 μM 3-isobutyl-1-methylxanthine, 0.2 mM indomethacin and 1 μM dexamethasone in final concentrations for 28 days. The medium was changed every other day. Differentiation was visualized by Oil Red O staining and confirmed with qPCR by testing mRNA level for fatty acid binding protein 4 (FABP4).

### Oil Red O staining

For visualizing lipid vesicles in the cells, the media was removed, and cells were gently washed with PBS. Cells were fixed in 10% formalin for 10-30 min. Formalin was removed, and wells were washed using PBS first and 60 % isopropanol last. Afterward, the wells were emptied to dry completely. Three parts of the Oil Red O stock solution (3% in isopropanol) were mixed with 2 parts of dH_2_O before staining to prepare the staining solution. After 10 min, the working solution was filtered and added to the dried wells for 10 min. The staining solution was removed, and cells were gently washed four times with dH_2_O and drying needed to be prevented. Staining was examined under the microscope and pictures were taken.

### Chondrogenic differentiation

Cells were seeded in a 24-well ultra-low attachment plate (10^6^ cells/well, Corning) in DMEM supplemented with 10 % FBS and 1% penicillin/streptomycin. During this cultivation step, the cells undergo mesenchymal condensation forming one self-organized aggregate. After condensation is completed (1-2 weeks), media was changed to chondrogenic conditions using serum-free DMEM (1 % penicillin/streptomycin) with 100 nM dexamethasone, 200 nM ascorbate phosphate, 40 μg/ml L-proline, 100 μg/ml sodium pyruvate, 1 % ITS-Premix and 10 ng/ml TGF-beta-3 (PromoKine) in final concentrations for 4 weeks. The media was changed 3 times a week. Differentiation was visualized after cryosectioning by proteoglycan staining with Alcian blue and Safranin O / Fast green on glass slides. The expression of collagen type II chain α1 (COL2A1) and aggrecan (ACAN) was further confirmed by qPCR and glycosaminoglycan content was determined.

### Cryosections

Cell condensates were embedded in O.C.T Compound (Tissue-Tek) and snap frozen and stored at −80 °C. Tissue was cut at −16 to −18 °C using a specialized knife for hard tissues and 8 μm sections were placed on glass slides (Histobond, Marienfeld, Germany) After drying, slides were stored at −20 °C. For RNA extraction samples were cut at a thickness of 25-30 μm and were collected in 2 ml centrifuge tubes. 1 ml QIAzol (Qiagen, Germany) was added, and RNA samples were stored at - 80 °C until extraction.

### Alcian blue staining

Sample slides were thawed at room temperature for 30 min and then fixed in 10% formalin for 10 min. After fixation was completed slides were brought to dH2O and then incubated in 3 % acetic acid for 3 min. Alcian blue staining using 1% Alcian blue 8G (Sigma Aldrich) in 3% acetic acid at pH 1.5-2.5 for 15-20 min was followed. Slides were washed in water than dehydrated and mounted. Glycosaminoglycans (GAG) were stained in turquoise to light blue.

### Safranin O / Fast green staining

Sample slides were thawed at room temperature for 30 min and then fixed in 10% formalin for 10 min. Slides were stained for 10 min in freshly mixed Weigert's iron hematoxylin solution (Roth). To differentiate the color slides were washed in running tap water for 10 min. Nuclear counterstain was performed by adding slides to Fast green solution (0.05% in dH_2_O) for 5 min. Sections were shortly (10 sec) rinsed with 1% acetic acid and then stained with Safranin O (0.1% in dH_2_O) for 5 min. Slides were dehydrated and mounted. Glycosaminoglycans were stained in red, nuclei in black and cytoplasm in light green.

### GAG measurement

The content of glycosaminoglycans (GAG) was measured using DMMB staining (1,9 dimethyl methylene blue). Therefore, condensates were first digested overnight at 56°C in 700 μl digestion buffer (50 mM Tris-HCl, 10 mM Na Cl, 3 mM MgCl2, 1% Triton X 100 at pH 7,9) containing proteinase K (50 ng/ml). The Digestion was stopped at 90 °C for 20 min. The sample was divided in half. One volume of 350 μl was used to isolate genomic DNA (Macherey Nagel, NucleoSpin Tissue Kit) the other half was further processed by digesting DNA with 3 units of DNase I at 37 °C overnight. The sample was centrifuged at 13,000 xg for 20 min, and the supernatant was divided into three samples of 100 μl each. 1 ml of DMBB solution (0.0016% DMBB, 5% EtOH, 0.2 M GuHCl, 0,2 % sodium formate and 0.2% formic acid, adjusted to pH 1.5 with HCl) was added to each 100 μl sample and were incubated for 30 min on shaker. The color complex was pelleted by centrifuging at 12,000 xg for 10 min. The supernatant was removed, and the pellet was resuspended on a shaker for 30 min by adding 300 μl DMBB de-complexation solution (4 M GuHCL in 50 mM sodium acetate (pH 6.8) + 10% 1-propanol). OD at 656 nm was measured in triplicates. Standard series of chondroitin sulfate in DMBB de-complexation solution (0-60 μg/ml) was performed for quantification. Column isolated DNA was quantified spectrometrically with the NanoDrop2000 (Thermo Scientific, USA) and glycosaminoglycan values were presented as ratio normalized to DNA content of related samples.

### RNA isolation

Adherent cells were lysed directly on the plate or after collection. Total RNA was extracted and purified using Nucleospin RNA II Kit (Macherey-Nagel, Germany) according to manufacturer protocol. For RNA isolation from cell condensates, samples were snap frozen and cut in 25-30 μm sections before lysis. Sections were transferred to 1 ml QIAzol (Qiagen) and stored at −80 °C until isolation. Phenol/chloroform extraction of total RNA was performed. RNA pellets were resuspended in 20-60 μl RNase free water depending on pellet size. RNA content was measured spectrometrically at 260 nm using NanoDrop2000 (Thermo Scientific, USA).

### Genomic DNA isolation

Genomic DNA was extracted with the NucleoSpin Tissue Kit (Macharey Nagel) according to the manufacturer’s protocol. DNA was eluted in 60 μl dH_2_O measured spectroscopically and stored at −20 °C.

Bisulfite sequencing of PECAM promoter

500 ng of genomic DNA was bisulfite converted using EZ DNA Methylation Gold Kit (Zymo Research Europe, Germany). The Sequence of the proximal promoter of PECAM was obtained from the UCSC genome browser (human genome assembly hg18). All cytosine bases not in a CpG dinucleotide context were converted to thymine in a text editor, and PCR primer were designed on the converted template using the primer3 software [19,20]. The PCR product was spanning the promoter from −164 bp to +285 bp regarding transcriptional start.

### Proliferation assay

Cells were harvested by trypsinization, washed twice with PBS and prepared in a cell density of 2 × 10^6^ cells/ml. A 2X CFDA-SE (10 μM) working solution was prepared. Equal volumes of cell suspension and the 2X working solution was mixed and incubated for 7 min at room temperature away from light. Labeling was stopped by adding 5 volumes of cold growth medium and incubation of 5 min on ice in the dark followed. Cells were centrifuged and washed twice with growth medium and then seeded in desired density to perform the assay. For the non-proliferating control cells were seeded at a high density to exclude cell proliferation by contact inhibition. Other wells were seeded at 20 % confluence to monitor cell doublings and were treated with 0.5 to 5 μM 5-aza-2’-deoxycytidine (Sigma Aldrich).

### cDNA synthesis and quantitative PCR

200 ng total RNA was reverse transcribed using the TaqMan kit (Thermofisher). The cDNA level was analyzed on MxPro3005 (Agilent Technologies) using SensiFast SYBR No-ROX Kit (Bioline, UK), 1 μl cDNA and 250 nM each primer. Quantitative PCR (qPCR) was done using the following thermal profile: 95 °C for 2 min, then 40 cycles of 12 sec at 95 °C, 7 sec at 64 °C and 3 sec at 72 °C with additional melt point analysis. Gene expression was calculated using an amplification efficiency (E) of 1.95 and values were normalized on reference gene UBE2D2 (NM_181838) expression.

### Statistics

For statistical analysis, the students t-test was performed and differences in expression called significant for p-values < 0.05 = * (< 0.01 = **, < 0.001 = ***, < 0.0001 = ****). Dependent on experimental set-up paired or un-paired t-tests were performed.

## Results

Primary chondrocytes de-differentiate upon isolation from articular cartilage characterized by a morphological transition from round to fibroblastoid shape. Interestingly, the impact on marker molecule expression seems to be bivalent (Fig 1). Although both tested isoforms of transcription factor SOX5 were down-regulated and the ECM molecule encoded by COL2A1 was not even detectable after cultivation, the mRNA levels for the cartilage-specific markers SOX6, SOX9, ACAN and PRG4 (proteoglycan 4) were unaffected after several weeks of cultivation.

**Figure 1:**
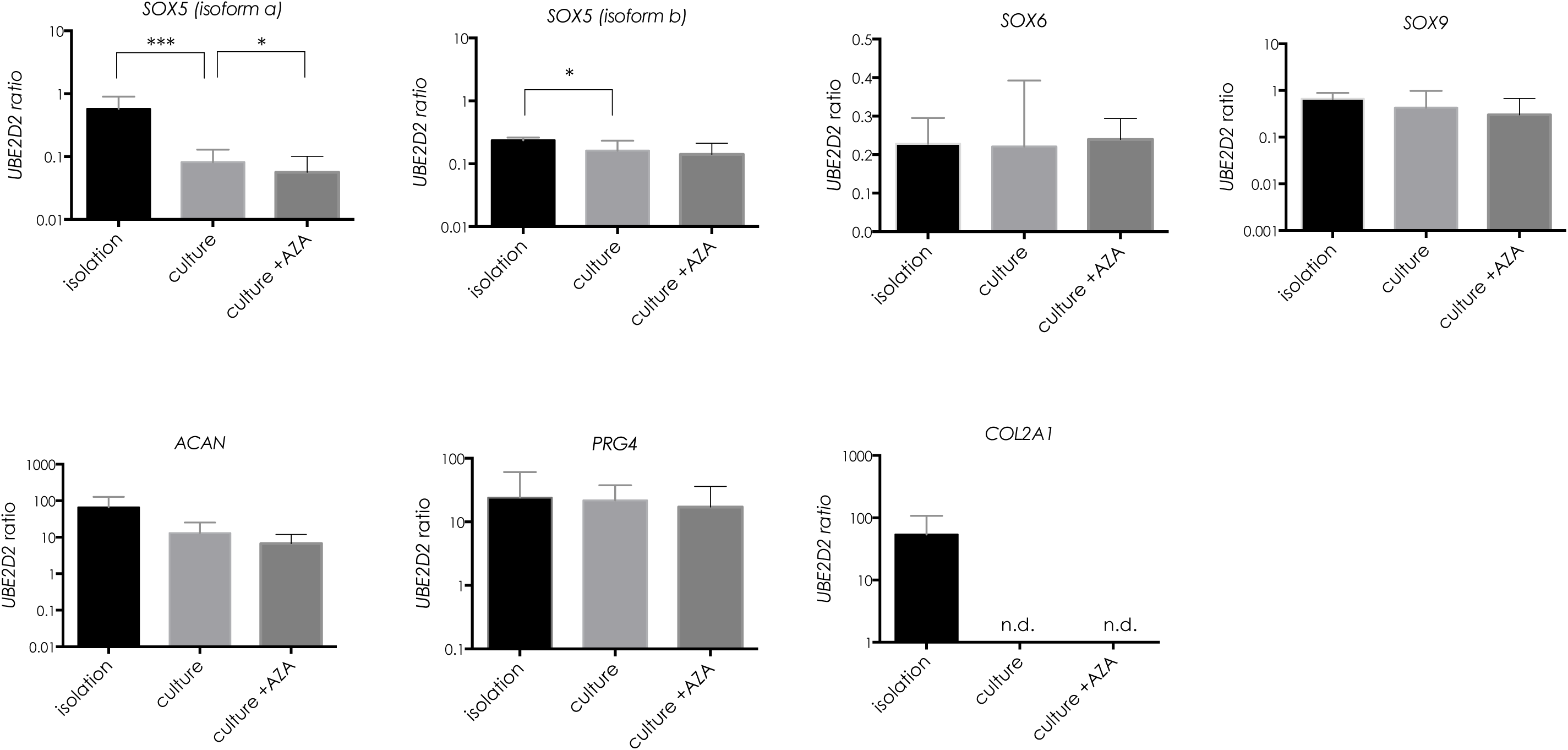
Gene expression in OA chondrocytes w/o 5-aza-dC treatment. mRNA levels of *SOX5*, *SOX6*, *SOX9*, *ACAN*, *PRG4* and *COL2A1* normalized on reference gene expression *UBE2D2*. Cells were cultivated for 4 passages and then treated with 1 μM 5-aza-dC for 4 weeks. Statistical analysis was done using ratio paired t-test. Values are the mean ± SD (n = 10; *p < 0.05, **p < 0.01, ***p < 0.001)

The impact of DNA methylation on marker gene expression was assessed by treating cultivated chondrocytes with the DNMT inhibitor 5-aza-2’-deoxycytidine. The treatment with 1 μM 5-aza-dC did not alter the expression of SOX5L (isoform b), SOX6, SOX9, PRG4, and ACAN. Only SOX5M (isoform a) was further down-regulated after 5-aza-dC administration. COL2A1 was not re-expressed after the treatment with the DNMT inhibitor (Fig 1).

Articular chondrocytes do not proliferate in their natural environment, but the cells re-enter the cell cycle after isolation and *in vitro* cultivation. Significant proliferation inhibition in chondrocytes was observed in our experiments with as little as 0.5 μM 5-aza-dC in the growth medium with no further gain of the effect between 2 and 5 μM 5-aza-dC after 8 days or 0.5 and 1 μM after 16 days of treatment (Fig 2).

**Figure 2:**
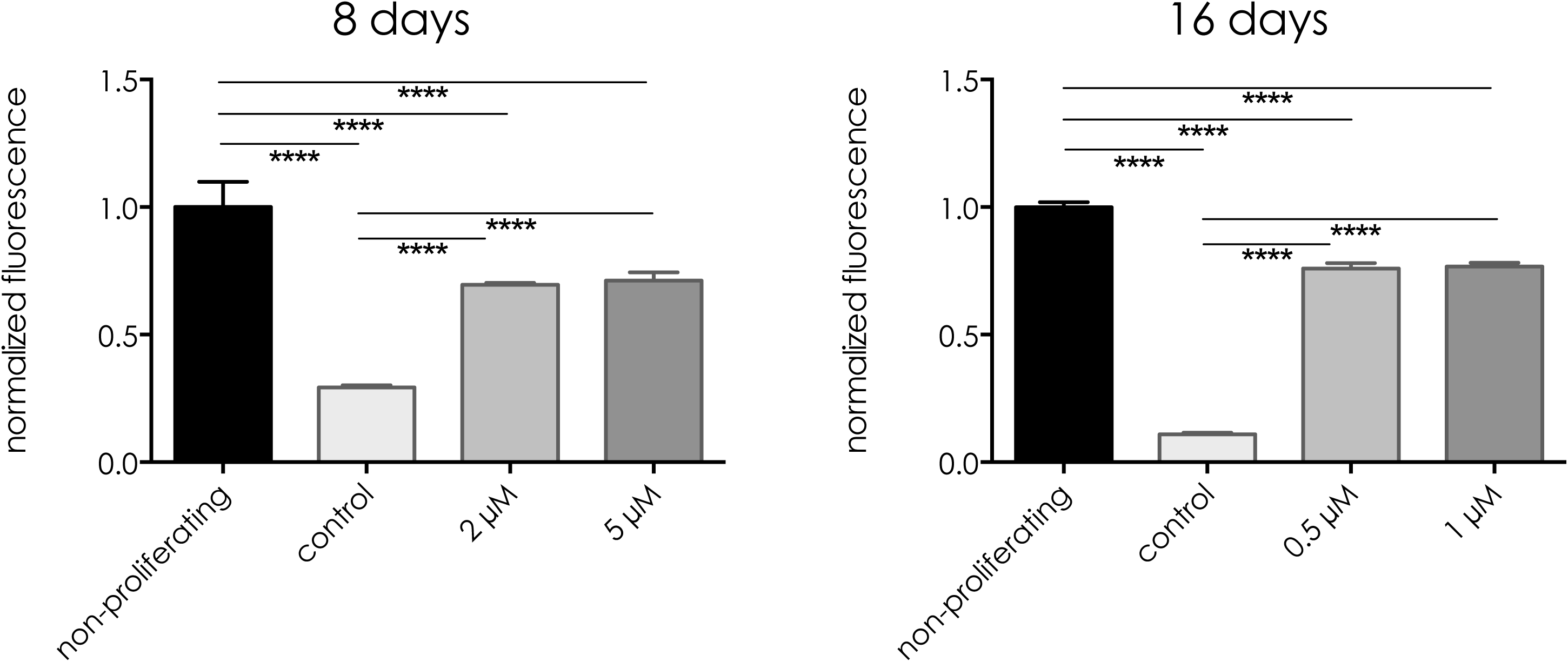
Inhibitory effect of 5-aza-dC on chondrocyte proliferation. CFSE labeled chondrocytes were treated with 2 or 5 μM 5-aza-dC for 8 days as well as 0.5 and 1 μM for 16 days. Values represent mean fluorescence (normalized on non-proliferative control). Statistical analysis was done using ratio paired t-test. Values are the mean ± SD (n = 10; *p < 0.05, **p < 0.01, ***p < 0.001)

Upon de-differentiation, the cells regain mesenchymal stromal cell properties like multiple differentiation potentials. To assess the influence of the DNMT inhibitor treatment, the capacity for adipogenic, osteogenic and chondrogenic differentiation was determined in the respective cells. Compared to the untreated control adipogenic and osteogenic differentiation was further improved by 5-aza-dC treatment (Fig 3 and 4). The staining of the calcified matrix by Alizarin Red as well as lipid vesicles by Oli Red O staining was positive for both populations after differentiation. However, DNMT inhibitor treated cells displayed a more pronounced lipid vesicle staining and higher expression of fatty acid binding protein-4 (FABP4). Furthermore, gene expression of osteopontin (OPN) showed increased levels after osteogenic differentiation in cells pre-treated with 5-aza-dC (Fig 4).

**Figure 3:**
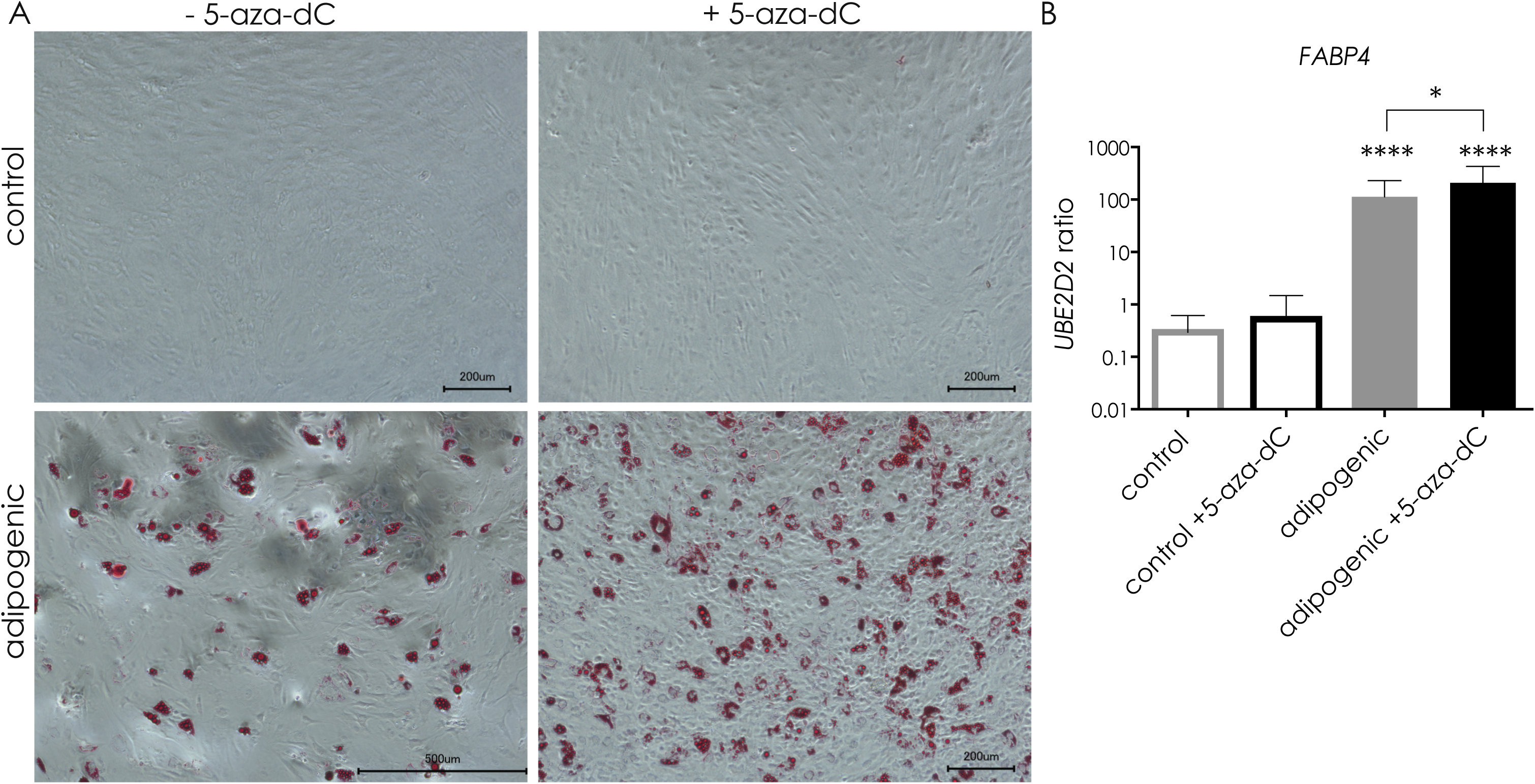
Influence of 5-aza-dC on the adipogenic potential of OA chondrocytes. After 5-aza-dC treatment, OA chondrocytes were differentiated towards adipocytes. (a) After 4 weeks, lipid vesicles were stained by Oil Red O. (b) Gene expression of fatty acid binding protein 4 (*FABP4*). Expression was normalized on reference gene *UBE2D2*. Values are mean ± SD and statistical analysis was done using paired ratio t-test. (n ≥ 4; *p < 0.05, ****p < 0.0001)

**Figure 4:**
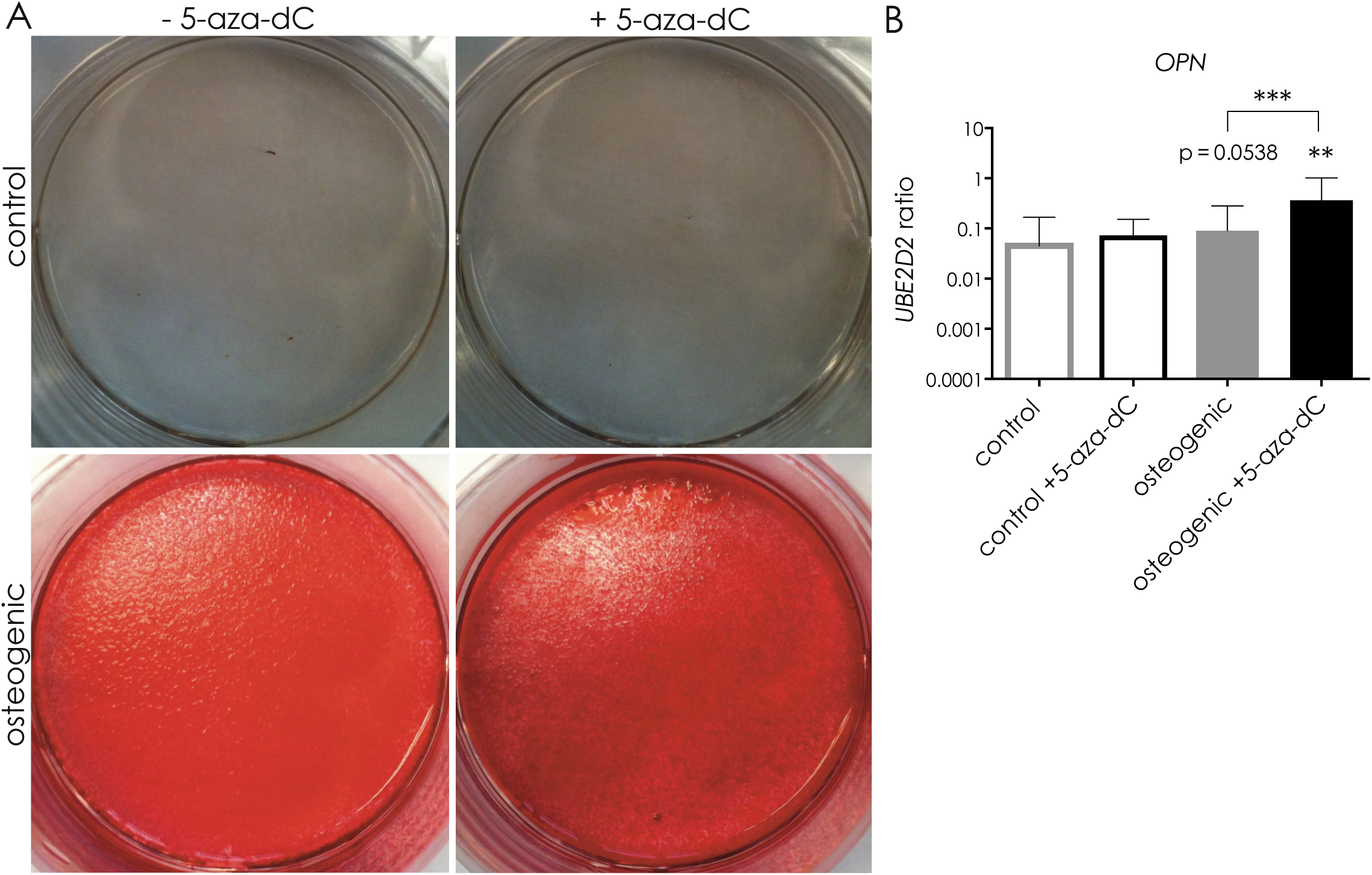
Influence of 5-aza-dC on the osteogenic potential of OA chondrocytes. After 5-aza-dC treatment, OA chondrocytes were differentiated towards osteoblast. (a) After 4 weeks, calcified matrix was stained by Alizarin Red. (b) Gene expression of osteopontin (*OPN*) was measured and normalized on reference gene *UBE2D2*. Values represent mean ± SD and statistical analysis was done using paired ratio t-test. (n ≥ 4; *p < 0.05, **p < 0.01, ***p < 0,001)

The most striking difference upon 5-aza-dC administration was observed during chondrocyte differentiation. Although no changes in cartilage marker expression were seen after 5-aza-dC treatment (Fig 1), the chondrogenic differentiation capacity was significantly compromised upon 5-aza-dC treatment (Fig 5). The stimulus of 3D culture and chondrogenic media was not sufficient to induce proper cartilage matrix formation (Fig. 5a), and glycosaminoglycan (GAG) content was significantly reduced (Fig 5b). Furthermore, the coupling of 5-aza-dC expansion with subsequent chondrogenic differentiation reduced the cartilage marker expression (Fig 6). These data strongly indicate that DNMT inhibition impaired a regulator of cartilage formation, whose activity is essential for active chondrogenic differentiation and assumedly dispensable for phenotype maintenance.

**Figure 5:**
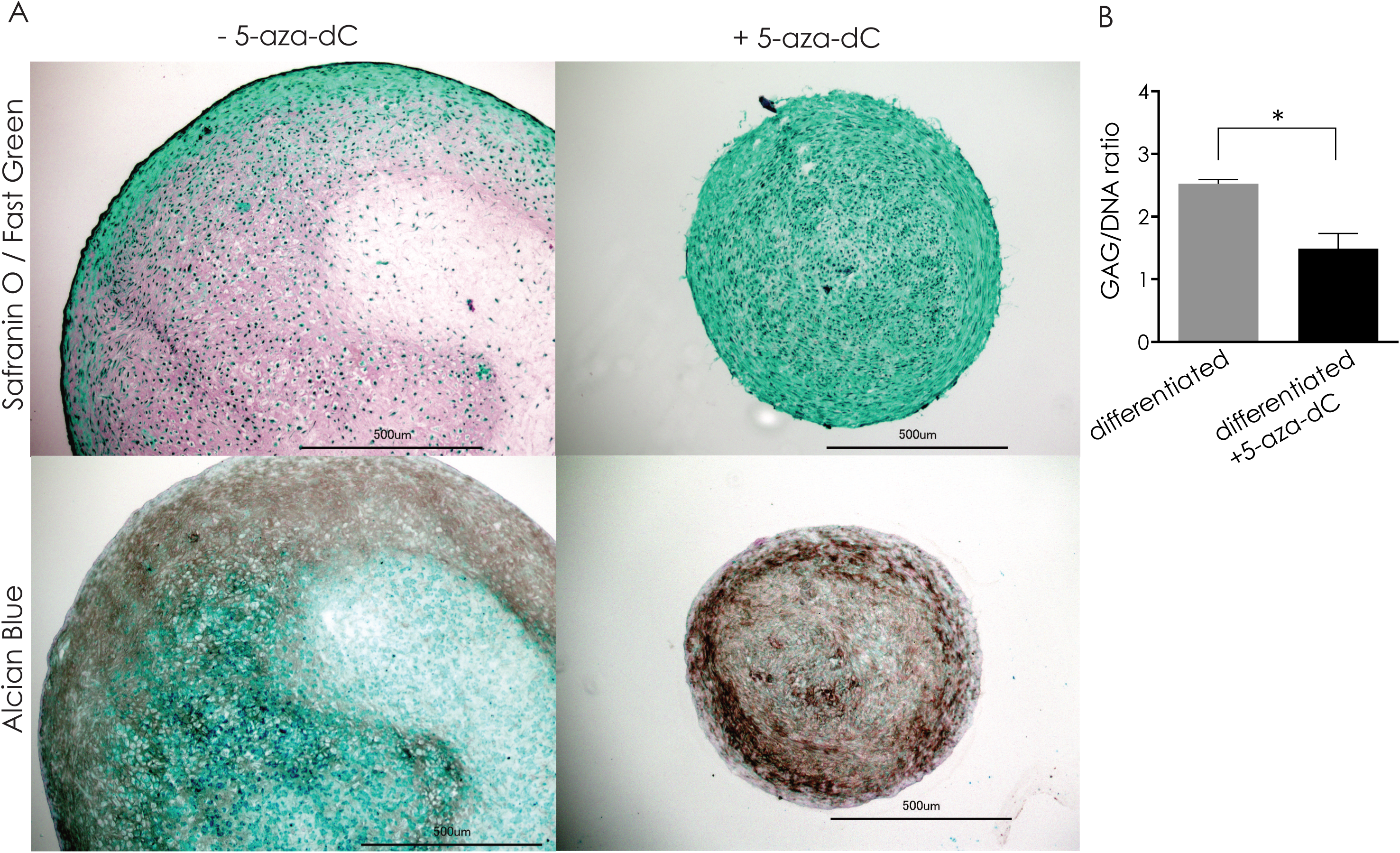
Effect of 5-aza-dC on cartilage matrix formation. After 5-aza-dC treatment, OA chondrocytes were differentiated towards cartilage for 6 weeks in 3D culture. (a) Frozen sections were stained for proteoglycans with Alcian Blue and Safranin O / Fast Green. (b) Glycosaminoglycan content was normalized on DNA amount of the same sample. Values represent mean ± SD and statistical analysis was done using paired ratio t-test. (n = 3; *p < 0.05)

**Figure 6:**
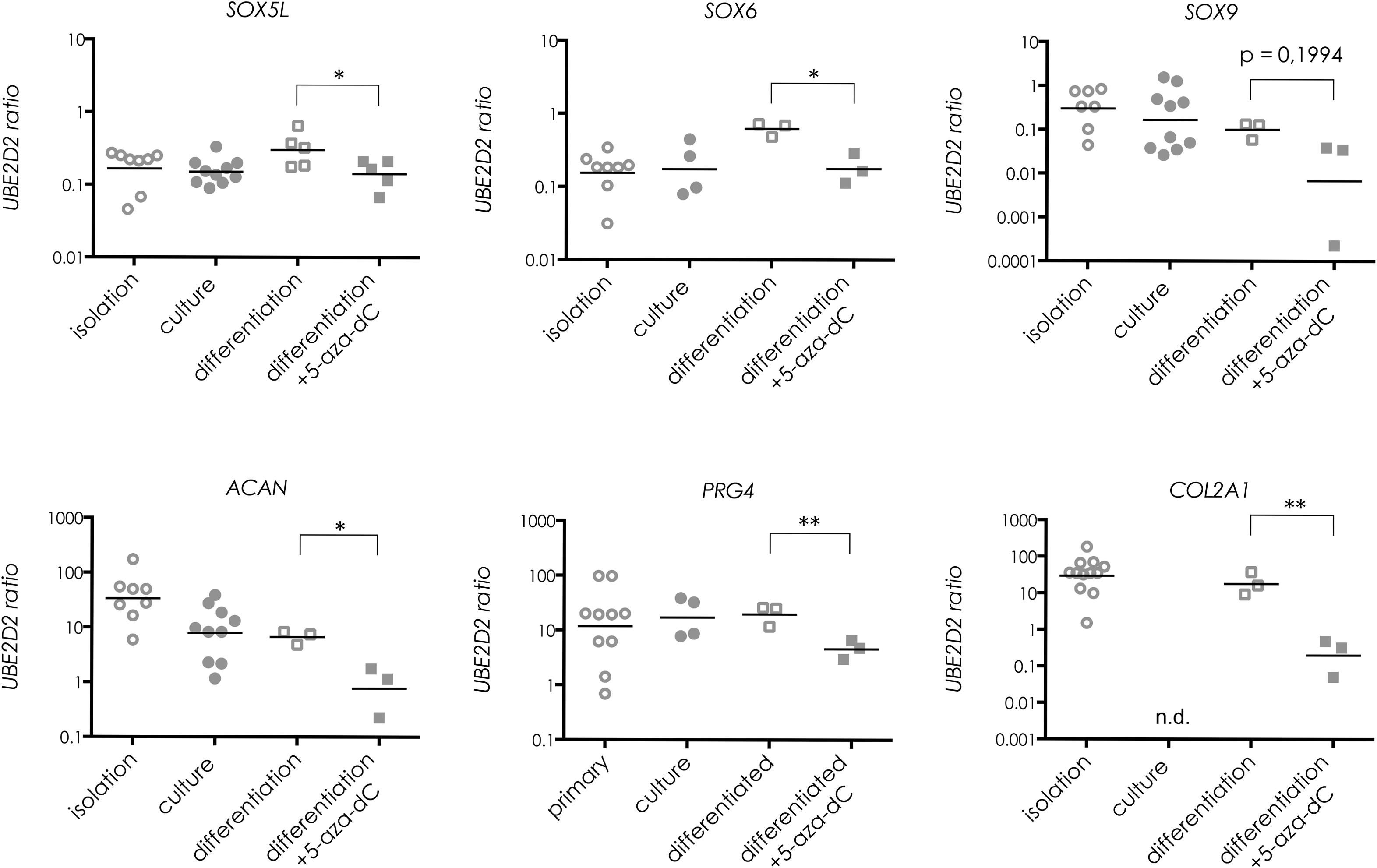
Influence of 5-aza-dC on cartilage marker expression. After 5-aza-dC treatment, OA chondrocytes were differentiated towards cartilage. After 6 weeks, gene expression of cartilage markers was measured and normalized on reference gene *UBE2D2*. Values represent mean ± SD and statistical analysis was done using paired ratio t-test. (n ≥ 4; *p < 0.05, **p < 0.01)

To assess the effect of DNMT inhibitor treatment on the DNA methylation status, the global DNA methylation level was determined comprehensively. As seen in Fig 7, even the long-term treatment of the isolated human OA chondrocytes with 1 μM 5-aza-dC exerted no significant influence on the global methylation level, while the treatment in HEK293T cells diminished DNA methylation after only 72 h. However, the analysis of an individual CpG at the site +242 of the PECAM promoter suggests that 5-aza-dC might act as a concentration-dependent DNMT inhibitor by de-methylation at the former strongly methylated sequence (Fig 7). To determine possible influences of cell culture or 5-aza-dC treatment on gene expression of DNA modifying enzymes, the gene expression of DNMTs were analyzed. The gene expression of DNMT1, DNMT3a, and DNMT3b was downregulated upon expansion *in vitro* with no further impact by additional DNMT inhibitor treatment (Fig 8).

**Figure 7:**
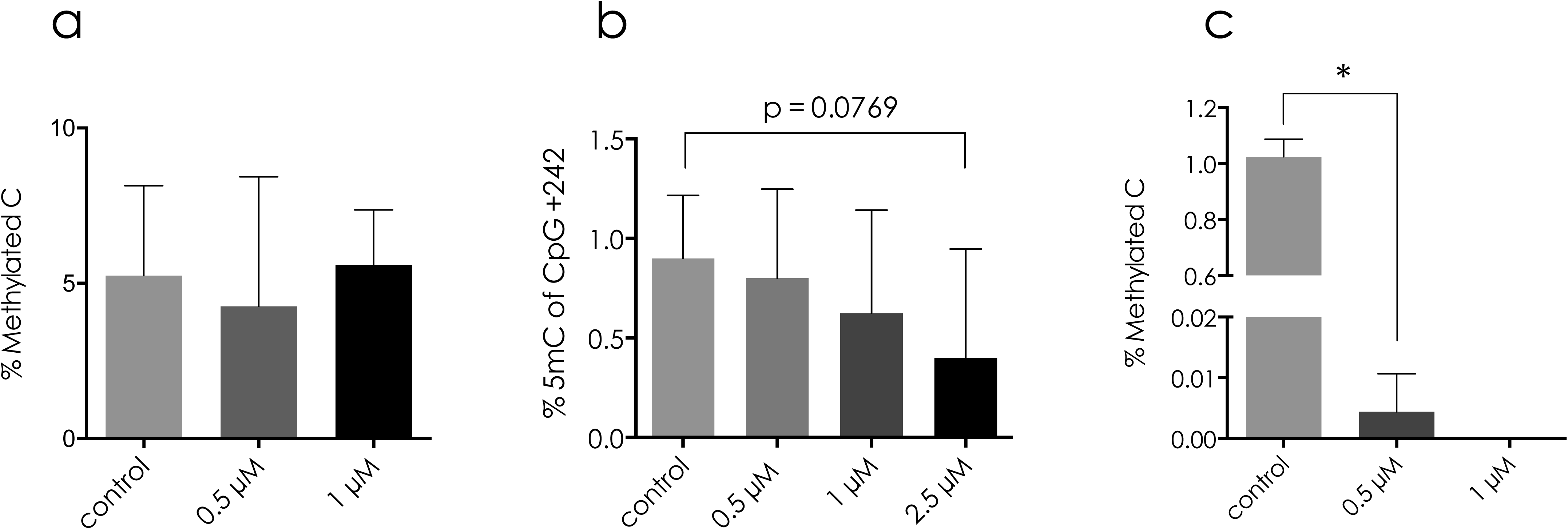
Global DNA methylation level in OA chondrocytes. (a) ELISA of 5mC in OA chondrocytes after 4 weeks of 5-aza-dC treatment. (b) Bisulfite sequencing of CpG at +242 of *PECAM* promoter after 4 week of 5-aza-dC exposure. (c) ELISA of 5mC in HEK293T cells after 72h of 5-aza-dC treatment. Values represent mean ± SD and statistical analysis was done using student t-test. (n = 4; *p < 0.05)

**Figure 8:**
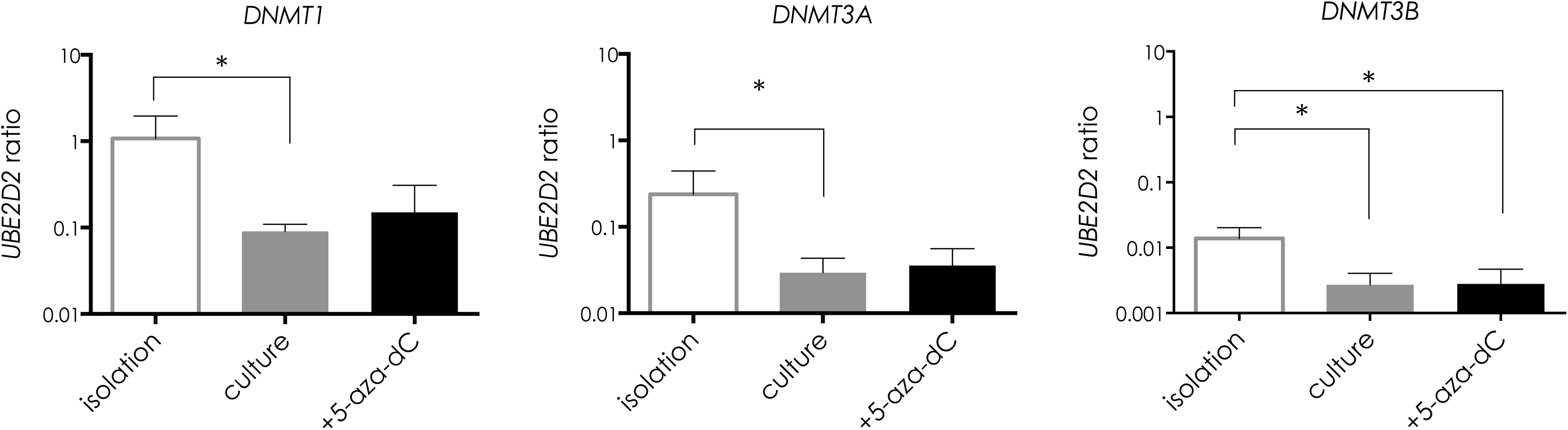
Gene expression of DNA methyltransferases *DNMT1*, *DNMT3A* and *DNMT3B.* Transcription level in isolated OA chondrocytes was compared with de-differentiated w/o 5-aza-dC treatment (4 weeks). Expression was normalized on reference gene *UBE2D2*. Statistical analysis was done using ratio paired t-test. Values represent mean ± SD (n ≥ 4; *p < 0,05)

## Discussion

Several publications report of de-differentiation and loss of functionality in healthy chondrocytes upon cell culture [21]. These studies mainly focus on the regenerative potential of chondrocytes. Even though the cartilage we extracted was from even and non-injured areas of the joint, the environmental signaling was nevertheless of late-stage OA and should be taken into consideration.

The transcriptional differences of healthy and osteoarthritic chondrocytes were described repeatedly. In these studies, the loss of cartilage marker expression was reported as shown in down-regulation of collagen type 2, aggrecan and SOX9 coupled with up-regulation of collagen type 1 [6,21]. Nevertheless, the de-differentiated chondrocytes showed a high potential for cartilage formation in 3D culture systems. As shown before, imitating the physiological environment of the tissue without further assistance was sufficient for the de-differentiated chondrocytes to re-differentiate [22]. We suggest that the chondrocytes of these OA patients had already experienced this transcriptional change. This could be the simplest explanation for the partial maintenance of marker expression upon cell culture, assuming the down-regulation already took place during disease progression (Fig 1). However, Lin et al. showed in 2008 similar gene expression profiles of cultured chondrocytes isolated from healthy and OA joints, indicating that after completed de-differentiation both chondrocyte populations were equal [23].

Comparative studies of healthy and OA chondrocytes revealed different DNA methylation pattern[24]. In our hands, treatment of OA chondrocytes with 5-aza-dC decreased cell proliferation already at low doses (Fig 2). In 2006, Unterberger et al. reported the subjacent mechanisms for the proliferation inhibition observed after DNMT reduction. The dissociation of DNMT1 from the replication fork activates a replication stress checkpoint in an ATR (ataxia telangiectasia mutated Rad3-related) dependent manner [25]. Through this mechanism one single incorporated cytosine analog could stall one replication complex until the missing enzyme is replaced, explaining the proliferative inhibition effect of small amounts of 5-aza-dC observed in our culture.

The increased expression of *OPN* and *FABP4* during osteogenic and adipogenic differentiation after 5-aza-dC treatment (Fig 3 and 4) are in line with the work of Kim et al., where the chondrogenic, osteogenic and neurogenic differentiation of human bone marrow MSCs was enhanced in the presence of 5-aza-dC [26]. In our hands, the most significant impact of 5-aza-dC treatment was observed on cartilage differentiation. Matrix production was reduced, and marker expression was down-regulated for the first time (Fig 5 and 6). Additionally, the expression of hypertrophic markers was elevated (Fig 7). These data are contrary to the work of Duan et al. in 2017, where the treatment of healthy chondrocytes with 5-aza-C could gradually reverse the de-differentiation of chondrocytes, shown by an increase of cartilage marker *SOX9* and the decrease of collagen type 1 expression thereby restoring the chondrogenic phenotype. Duan et al. isolated chondrocytes from trauma patients while chondrocytes in this work were extracted from late-stage OA joints. Duan et al. used 2 μM 5-aza-C (RNA nucleotide) for 24 hours, while in our lab hands cells were treated for 3 to 4 weeks with 1 μM 5-aza-dC (DNA nucleotide). Komashko and Fanham reported the differences between long- and short-term treatment of 5-aza-C on DNA methylation and histone modification patterns in 2010. They showed a regulatory effect of the inhibitor mostly on genes which were already unmethylated before treatment and that short-term treatment only slightly reduced the DNA methylation of HEK293 cells. Alvarez-Garcia et al. analyzed in 2016 the influence of short (48 h) and long-term (4-5 weeks) treatment of 5-aza-dC by determining the expression of few tested markers in chondrocytes, where short-term treatment enhanced the transcription of more genes compared to long-term stimulus. Epigenetic differences between normal and osteoarthritic chondrocytes are described by Alvarez-Garcia et al., where the DNA methylation profiles significantly differ between these two groups [24]. Taken these findings together, the duration of treatment influences transcription by different mechanisms, since in short-term treatment the incorporation rate is very low due to the few numbers of cell doublings during that time frame.

The long-term treatment of OA chondrocytes with 5-aza-dC in our lab did not affect the global DNA-methylation levels (Fig 8). In the work of Hashimoto et al. the incubation of 5-aza-dC in chondrocytes led to a decrease of DNA methylation of specific CpGs within the IL1b promoter while global DNA demethylation was not shown [28]. In 2017, a methylome analysis by NGS performed by Duan et al. illustrated the differences in DNA methylations of chondrocytes at different passages as well as 5-aza-dC treated versus untreated cells. Their findings indicate a directed change in DNA methylation, since most prominently the regions which were hypomethylated by 5-aza-dC, got hypermethylated upon artificial cell culture. Furthermore, the treatment of 5-aza-dC in primary healthy chondrocytes indeed de-methylate DNA, but preferably of distinct and possibly cartilage-specific regions. This was also published for human cancer cell lines by Hagemann et al., where the DNA de-methylation was found to be non-randomly and reproducible. They reported that upon treatment with 5-aza-dC specific CpGs within CpG islands become re-methylated and furthermore they identified sequences which were never affected by treatment at all [29]. The inhibiting effect of 5-aza-dC on cartilage marker expression during chondrogenesis is the focus of further investigations. Whole genome methylation analysis is required to identify the key player of the observed changes with regards to the duration of treatment.

Without fully exploring the mechanisms behind the cartilage marker maintenance in OA chondrocytes, their beneficial properties represent a helpful tool for *in vitro* applications and our data are in line with multiple publications, which report that OA chondrocytes are a source of osteochondroprogenitors and have equal cartilage formation capacity as healthy chondrocytes [30–33].

## Conclusion

Chondrocytes isolated from late-stage osteoarthritis do not show the same changes as healthy chondrocytes during de-differentiation upon cell culture. They can differentiate towards adipogenic, osteogenic and chondrogenic lineages. The adipogenic and osteogenic differentiation was further enhanced under long-term treatment with DNMT inhibitor 5-aza-dC, while the proliferation of cells slowed down. On the other hand, the chondrogenic differentiation was inhibited. Although the DNMT inhibitor was added every second day for four weeks during the expansion of cells, no global DNA de-methylation could be observed, indicating an additional independent mechanism responsible for the observed effects.

## Supporting information

## List of abbreviations

5-aza-C: 5-aza-cytidine
5-aza-dC: 5-aza-2’-deoxycytidine
5mC: 5-methyl-cytosine
ACAN: aggrecan
COL2A1: collagen type II chain alpha 1
DNMT: DNA methyltransferase
FABP: fatty acid binding protein
GAG: glycosaminoglycan
NGS: Next generation sequencing
OA: osteoarthritis
OPN: osteopontin
PRG4: proteoglycan-4
UBE2D2: ubiquitin-conjugating enzyme E2 D2

## Declarations

### Ethics approval and consent to participate

The acquisition of cartilage tissue was approved by the Ethics Committee Charité and all patients gave written consent for this research.

### Consent for publication

Not applicable.

### Availability of data and materials

All data generated or analyzed during this study are included in this published article and its supplementary information files.

### Authors’ contributions

SK designed the experimental setting, performed cell culture and analyzed the results. LRS and OV performed parts of the cell culture as well as the molecular biological methods. All authors read and approved the final manuscript.

## Acknowledgments

We acknowledge support by the German Research Foundation and the Open Access Publication Funds of TU Berlin

