## Supplementary material for "Effects of 5-aza-2’-deoxycytidine on human osteoarthritic chondrocytes"

Table 1 (supplement)

| target | primer up (5´-3´) | primer down (5´-3´) |
|---|---|---|
| *ACAN* | GGGATGGTGGATGTCAGTTGG | CTCCTGCCTCTTGGGCTGTT |
| *COL2A1* | ACTCAAGTCCCTCAACAACCAG | CTGCTCCACCAGTTCTTCTTG |
| *DNMT1* | GGTGGTGGATGACAAGAAGTTTG | TGAGGATGGGCTGGTACTGTG |
| *DNMT3A* | TCCAACCCTGTGATGATTGATG | CTTTGCTGAACTTGGCTATCCTG |
| *DNMT3B* | CCGGTGTTTCTGTGTGGAGTG | CTTCATATTCAAGCCCCGTGTC |
| *FABP4* | GCAGAAATGGGATGGAAAATCA | CGTCCCTTGGCTTATGCTCTC |
| *OPN* | CACTGATTTTCCCACGGACCT | CCATTCAACTCCTCGCTTTCC |
| *PRG4* | CCATGCTTTCCGATGAGACC | CAATGGGGGAAGGAATACCC |
| *SOX5 isoform b* | CGAGCCACCAAAACCCATC | TCAGCAAGAGGAAAGCCCAGTAG |
| *SOX5 isoform a* | TTGACAGGTTCAGTTGGAGACG | GAGTGAGGCTTGTTGGGAAAAC |
| *SOX6* | TAAATACAAACCCCGACCGAAAC | GATAGCACCAGGATACACAACACCT |
| *SOX9* | GCCAGGTGCTCAAAGGCTAC | CGCTTCTCGCTCTCGTTCA |
| *UBE2D2* | TCAGGCACTAAAGGATCATCTGG | TCTTGACAATTCATTTCCCAACAG |

Table 1: Sequences of qPCR primer
